# Fate-mapping within human kidney organoids reveals conserved mammalian nephron progenitor lineage relationships

**DOI:** 10.1101/432161

**Authors:** Sara E Howden, Jessica M Vanslambrouck, Sean B Wilson, Ker Sin Tan, Melissa H Little

## Abstract

While mammalian kidney morphogenesis has been well documented, human kidney development is poorly understood. Here we combine reprogramming, CRISPR/Cas9 gene-editing and organoid technologies to study human nephron lineage relationships *in vitro*. Early kidney organoids contained a *SIX2*^+^ population with a transcriptional profile akin to human nephron progenitors. Lineage-tracing using gene-edited induced pluripotent stem cell (iPSC) lines revealed that *SIX2*-expressing cells contribute to nephron formation but not to the putative collecting duct epithelium. However, Cre-mediated temporal induction of the *SIX2*^+^ lineage revealed a declining capacity for these cells to contribute to nephron formation over time. This suggests human kidney organoids, unlike the developing kidney *in vivo*, lack a nephron progenitor niche capable of both self-renewal and ongoing nephrogenesis. Nonetheless, human iPSC-derived kidney tissue maintains previously identified lineage relationships supporting the utility of pluripotent stem cell-derived kidney organoids for interrogating the molecular and cellular basis of early human development.

## Introduction

Our understanding of kidney morphogenesis is predominantly based on studies performed in model organisms. Gene-knockout and fate-mapping studies, performed largely in mouse, have uncovered many of the fundamental molecular processes underlying kidney development, homeostasis, and disease (Humphreys and DiRocco, 2014). While inferences can be made using a well-characterized mammalian model, our capacity to validate the true relevance of lineage relationships or key genetic pathways in human development is hampered by the scarcity of human fetal tissue for experimental purposes. Recent analyses of human fetal kidney has provided important insights and highlighted several differences between human and mouse at both the immunohistochemical and transcriptional level (Lindstrom et al., 2018a; Lindstrom et al., 2018b; O’Brien et al., 2016). However, the examination of human fetal tissue is ethically constrained, provides only snapshots at a fixed developmental time-point, and does not represent an ideal platform for evaluating whether differences between human and model organisms convey functional relevance.

The capacity to create a model of the developing kidney *in vitro* provides a unique opportunity to better understand nephrogenesis at the molecular level in a human context. Several protocols have now been described for the directed differentiation of human pluripotent stem cells (hPSCs) to kidney organoids (Morizane et al., 2017). We have established a protocol that generates complex multicellular kidney organoids containing patterning and segmenting nephrons, endothelial, perivascular and stromal cells (Takasato et al., 2016; Takasato et al., 2015). After day 25 of differentiation, a single organoid contains up to 100 nephrons, each beginning to show functional maturation with podocyte foot process formation and albumin uptake in proximal tubules. Moreover, kidney organoids exhibit a transcriptional profile that is remarkably similar to first trimester human fetal kidney (Takasato et al., 2015). Combined with an ability to generate developing kidney tissue *in vitro*, improvements in the speed and accuracy with which it is possible to edit the genome of hPSCs using CRISPR/Cas9 technology (Howden et al., 2018) provides an unprecedented opportunity to interrogate the molecular and cellular basis of early human development using hPSC-derived human tissue.

During kidney morphogenesis in the mouse, new nephrons form throughout development from a multipotent self-renewing mesenchymal population located at the periphery of the developing kidney (Boyle et al., 2008; Kobayashi et al., 2008). These nephron progenitor cells (NPCs), marked by expression of *Six2* and *Cited1*, exist in close association with the tips of the branching ureteric epithelium. Nephron formation involves a mesenchyme to epithelial transition to form a renal vesicle with this process being accompanied by downregulation of *Six2* (Little and McMahon, 2012; Short et al., 2014). Although *Six2* expressing cells are capable of giving rise to all cell types of the nephron (Kobayashi et al., 2008), *Six2*-expressing NPCs are not the source of the branching ureteric epithelium (Boyle et al., 2008; Kobayashi et al., 2008), which is instead derived from a distinct ureteric progenitor population (Taguchi et al., 2014).

In this study, we use numerous genetically engineered human induced pluripotent stem cell (iPSC) lines to interrogate nephron formation during kidney organoid differentiation. A *SIX2* reporter line was generated to monitor potential NPCs during kidney organoid differentiation, with reporter gene expression detected early after organoid formation. Moreover, lineage-tracing experiments using iPSCs harboring Cre recombinase within the endogenous *SIX2* locus, in addition to a ubiquitously expressed loxP-flanked fluorescence cassette, demonstrate that these *SIX2*-expressing cells can contribute to nephron formation. While *SIX2*-derived cells formed proximal nephron segments, they were absent from the GATA3^+^CDH1^+^ epithelium, consistent with a collecting duct identity for the latter. These findings illustrate the feasibility of combining genome engineering, stem cells and organoid technologies to interrogate and dissect human lineage relationships *in vitro*.

## Results

### SIX2 Expression Persists Throughout Kidney Organoid Differentiation

Previous studies established *Six2* as a specific marker of NPCs in the developing mouse kidney (Kobayashi et al., 2008; Self et al., 2006). More recently, immunohistochemical and transcriptional analyses of human fetal kidney suggest *SIX2* expression also marks NPCs in the developing human kidney (Lindstrom et al., 2018a; O’Brien et al., 2016). To interrogate the presence and potential of NPCs during the course of human kidney organoid differentiation, we generated *SIX2* fluorescent reporter iPSCs using CRISPR/Cas9 to knock-in an EGFP cassette under the transcriptional control of the endogenous *SIX2* locus (Figure 1A). Several clonally derived iPSC lines with either heterozygous or homozygous insertion of the EGFP reporter were established using a previously described method that combines reprogramming and gene editing together in a single step (Howden et al., 2015; Howden et al., 2018). Both heterozygous (SIX2^EGFP/+^) and homozygous (SIX2^EGFP/EGFP^) clones, distinguished by PCR analysis (Supplementary Figure 1A), were used in subsequent differentiation experiments. Kidney organoids were generated using our previously described protocol (Takasato et al., 2016) (Figure 1B) with two modifications: 1) TeSR-E6 was used instead of APEL as the base differentiation medium and 2) a 3D Bioprinter was used to transfer day 7 aggregates to Transwell filters instead of manual transfer with a handheld pipette. This modified protocol enables the generation of large numbers of highly reproducible organoids that are equivalent at the level of morphology, component cell types and gene expression to those previously reported via manual generation (Higgins et al., unpublished data). Reporter gene expression in SIX2^EGFP^ organoids was monitored routinely by flow cytometry and fluorescent microscopy and was first detected at approximately day 10 of differentiation, consistent with previous reports (Morizane et al., 2015; Taguchi et al., 2014; Takasato et al., 2015). Notably, reporter gene expression was maintained until the cessation of differentiation at day 25 (Figure 1C). Co-localization of EGFP and SIX2 was confirmed by RT-PCR analysis of sorted EGFP-expressing and non-expressing fractions (Supplementary Figure 1B) and by immunofluorescence (Figure 1D), where SIX2 was restricted to the interstitial/mesenchymal compartment (Supplementary Figure 1C). Organoids derived from either SIX2^EGFP/+^ or SIX2^EGFP/EGFP^ iPSCs also exhibited highly similar dynamics with respect to reporter gene expression and nephron formation, although reporter gene intensity was greater in SIX2^EGFP/EGFP^ kidney organoids as expected (data not shown). Reporter expression was also dependent on CHIR99021 concentration and duration during the first stage of differentiation (Supplementary Figure 1D), consistent with previous studies showing that increased WNT signaling during hPSC differentiation induces a more posterior intermediate mesoderm (Takasato et al., 2015). Organoids generated from cultures treated with < 4 μM CHIR99021 failed to induce SIX2 expression or any recognizable epithelial structures, whereas differentiations performed with 8 μM CHIR for > 4 days contained the highest fraction of *SIX2*-expressing cells (>50%). Cultures treated with 6–8 μM CHIR99021 for 4 days formed organoids with the most balanced profile with respect to relative abundance of NEPHRIN^+^ podocytes, LTL^+^ proximal tubules, ECAD^+^ distal tubule structures, and putative GATA3^+^/ECAD^+^ collecting duct epithelium, as determined by whole-mount immunostaining (Supplementary Figure 1E). This condition was used for all subsequent differentiations.

**Figure 1.**
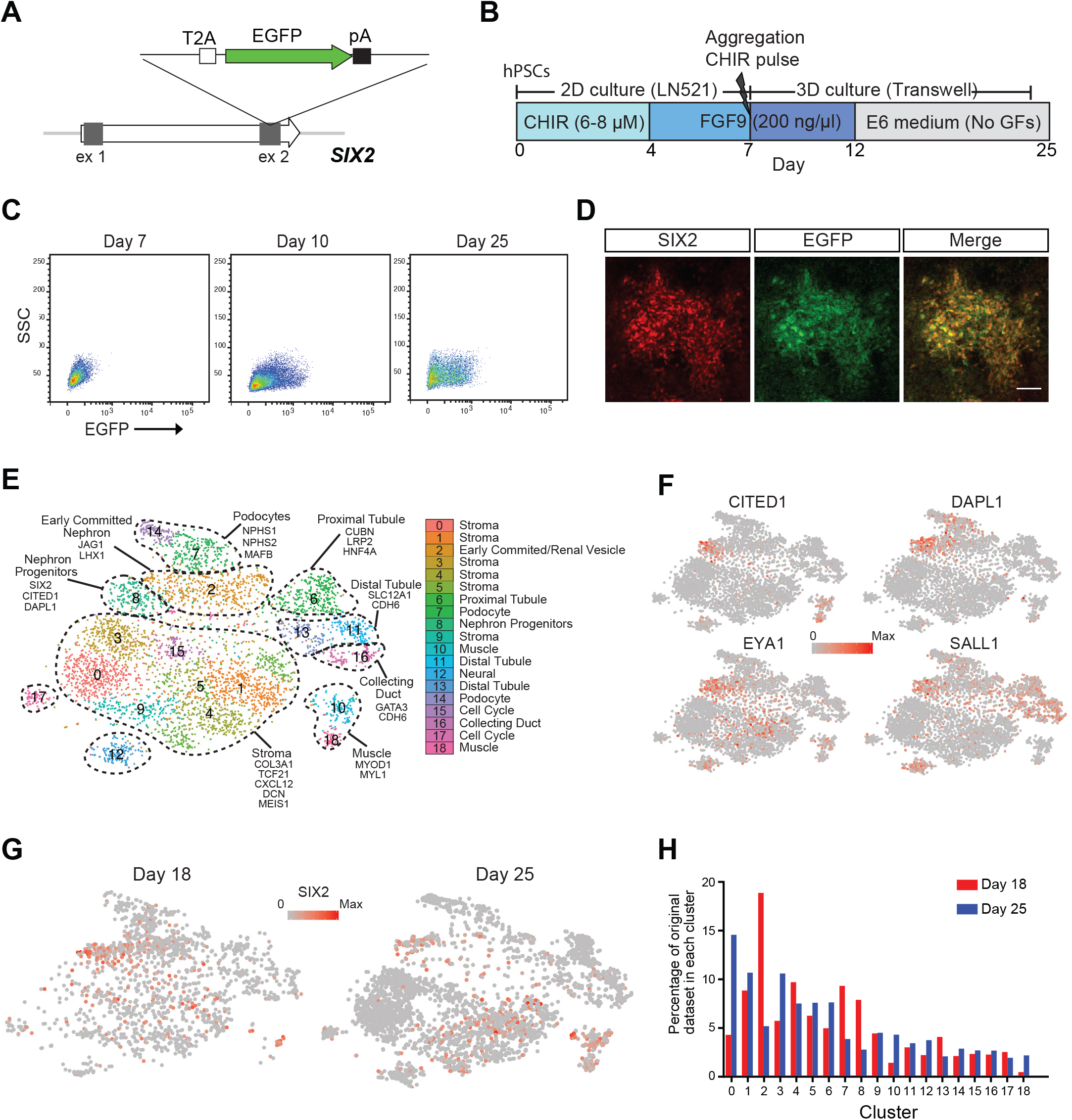
Analysis of *SIX2*-expressing Cells during Kidney Organoid Development. (A) Schematic diagram of the targeting strategy for generation of SIX2 reporter iPSCs. The EGFP gene was inserted just upstream of the *SIX2* stop codon, linked via the self-cleaving T2A sequence. (B) Outline of the kidney differentiation protocol used throughout this study. (C) Flow cytometry analysis of kidney organoids derived from SIX2^EGFP/EGFP^ iPSCs. (D) Immunostaining confirming co-localization of SIX2 and EGFP in SIX2^EGFP/EGFP^ kidney organoids. (E) tSNE plot representing the merged datasets from single cell RNAseq analysis of day 18 (1865 cells) and day 25 (3500 cells) organoids. Nineteen clusters were identified. (F) A distinct SIX2-expressing cluster was identified as a nephron progenitor-like population based on expression of other known markers, including *CITED1*, *DAPL1*, *EYA1* and *SALL1*. (G) *SIX2* marks a diverse range of cell types within human kidney organoids with *SIX2* transcripts detected in numerous clusters in both day 18 and day 25 organoids. (H) Graphical representation showing the cell populations that are enriched or depleted in day 18 versus day 25 kidney organoids.

### Single Cell RNAseq of Whole Kidney Organoids Identifies Multiple SIX2-expressing populations

To examine *SIX2*-expressing cells in greater depth during the process of kidney organoid differentiation, single cell transcriptome profiling of day 18 and 25 organoids was performed using the 10X Chromium platform. Nineteen cell populations emerged from guided clustering analyses using Seurat (Butler et al., 2018), with several of these pertaining to different nephron segments and cells at various stages of nephrogenesis (Figure 1E). Multiple stromal populations were also identified, which expressed collagens *COL1A1* and *COL3A1*, as well as kidney stromal markers *DCN* and *CXCL12*. With respect to *SIX2*-expressing cells, a distinct population (cluster 8) exhibited strong congruence with human fetal nephron progenitors, as determined by co-expression of several other previously described NPC markers, including *CITED1*, *DAPLI*, *EYA1* and *SALL1* (Lindstrom et al., 2018a; Morizane et al., 2015; Taguchi et al., 2014) (Figure 1F). Notably, *SIX2* expression was not solely restricted to putative NPCs, with *SIX2* transcripts detected in several additional clusters, including a subset of the renal stroma, and within an “off-target” population that displayed a muscle-like transcriptional profile (Figure 1G). This *SIX2-* expressing muscle-like population was clearly evident in day 25 organoids, but largely absent in early organoids (Figure 1G and Figure 1H). Compared to day 18, day 25 organoids also showed an overall reduction in both the *SIX2*-expressing NPCs and ‘early committed nephron’ clusters (Figure 1H). Taken together, these findings demonstrate that *SIX2* marks several distinct cell types within human kidney organoids, including a putative NPC population, which is enriched in early kidney organoids but depleted as differentiation proceeds. It also suggests that nephrogenesis is a transient process within organoids with evidence for nephron maturation with time.

### Generation of a Dual Fluorescence Cassette for Human Fate-Mapping Studies

To facilitate fate-mapping experiments in iPSC-derived kidney organoids and ultimately determine whether *SIX2*-expressing cells can give rise to nephrons, we generated a dual fluorescence cassette comprising a loxP-flanked EGFP and adjacent mCherry reporter for incorporation into the endogenous *GAPDH* locus (Figure 2A). We have previously shown that this locus facilitates ubiquitous and consistent transgene expression in hPSCs, both prior to and following differentiation into various different cell types (Kao et al., 2016). The functionality of our fluorescence cassette was validated following introduction into the human embryonic stem cell line, H9, where correctly targeted cells could clearly be identified by expression of the EGFP reporter (Figure 1A). EGFP-expressing clones were isolated, expanded and subsequently transfected with mRNA encoding Cre recombinase. This resulted in the rapid induction of mCherry expression and a corresponding loss of EGFP expression within 8 hours post-transfection in > 95% of cells (Figure 2B). Kidney organoids were successfully derived from EGFP or mCherry-expressing hPSCs, representing cells before and after exposure to Crerecombinase respectively, where we observed maintenance of appropriate reporter gene expression in all component cell types as determined by live fluorescent microscopy and flow cytometry (Figure 2C and D).

**Figure 2.**
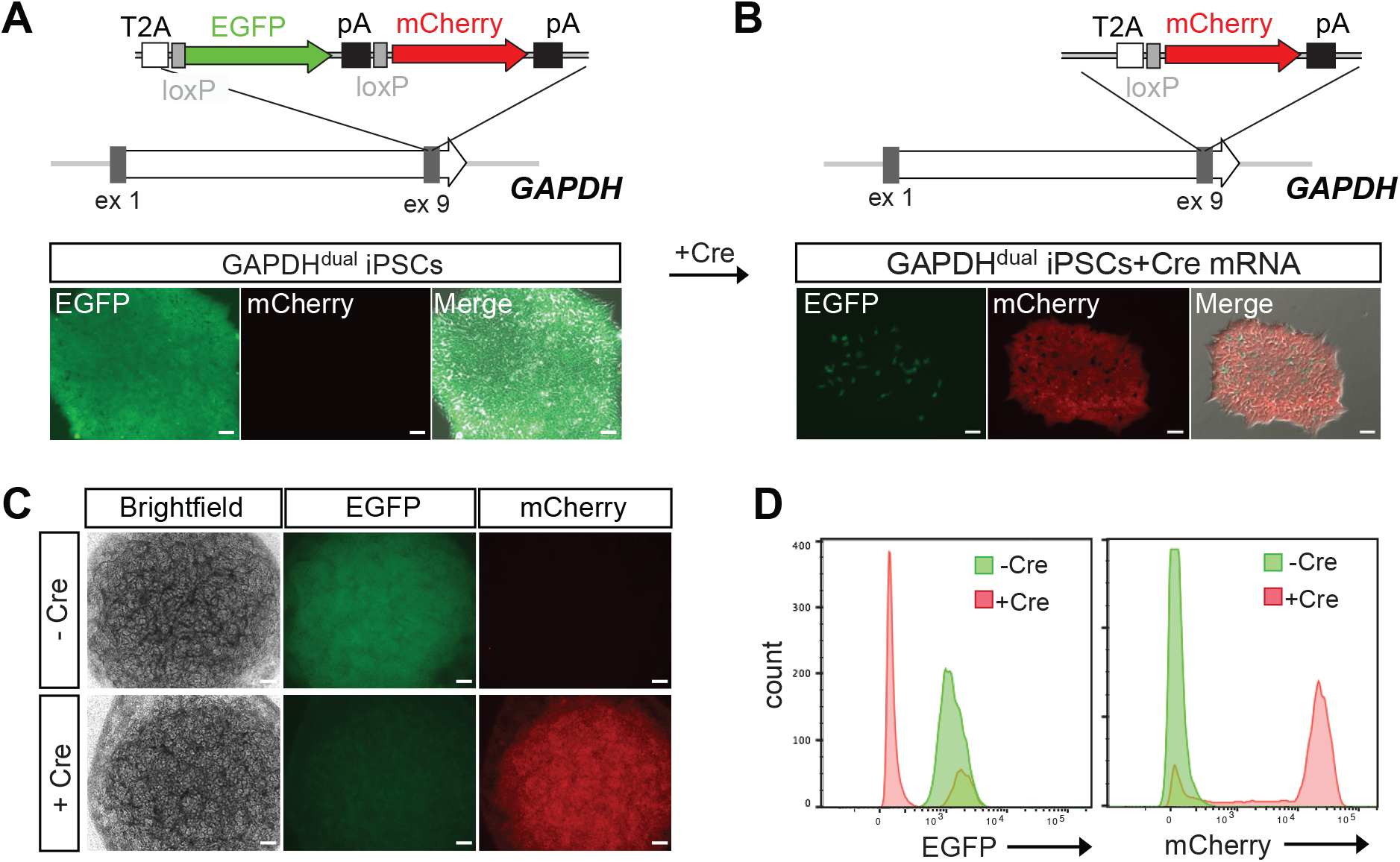
Generation and Characterization of a Dual Fluorescent Reporter Construct for Downstream Lineage Tracing Experiments. (A) Schematic diagram of the targeting strategy. A loxP-flanked EGFP and adjacent mCherry reporter were inserted downstream of the endogenous *GAPDH* coding region, linked via a self-cleaving T2A sequence. In the absence of Cre recombinase, cells constitutively express EGFP. (B) The loxP-flanked EGFP gene is deleted following Cre-mediated recombination. Shortly after (< 8 hours) the transient transfection of mRNA encoding Cre recombinase, GAPDH^dual^ hPSCs permanently switch from EGFP to mCherry reporter gene expression. Reporter gene expression is maintained in all cell types within kidney organoids generated from GAPDH^dual^ hPSCs before and after exposure to Cre, as detected by fluorescent microscopy (C) and flow cytometry (D). Scale bar = 50 μm (panels A and B) and 200 μm (panel C).

### SIX2^+^ Cells Contribute to Nephrons in Human Kidney Organoids

Previous fate-mapping studies performed in mouse demonstrated that *Six2* expression marks a multipotent nephron progenitor population that gives rise to all cell types of the of the nephron, from the glomerular epithelial populations through to the connecting segment (Georgas et al., 2009; Kobayashi et al., 2008). We decided to perform a similar fate-mapping analysis to examine whether *SIX2*-expressing cells can contribute to nephron formation in hPSC-derived kidney organoids. Clonally derived iPSCs with homozygous insertion of the Cre recombinase gene at the 3’ end of the *SIX2* coding region were established using our one-step reprogramming/gene-editing protocol (Howden et al., 2018) and were subsequently used for insertion of the dual fluorescence cassette into the *GAPDH* locus (Figure 3A). EGFP-expressing colonies, hereafter referred to as SIX2^Cre/Cre^:GAPDH^dual^ iPSCs, were identified by fluorescent microscopy, isolated and expanded for downstream differentiation experiments. Flow cytometry analysis of kidney organoids generated from SIX2^Cre/Cre^:GAPDH^dual^ iPSCs showed mCherry^+^ cells beginning to emerge at approximately day 10 of differentiation, coinciding with activation of endogenous *SIX2* and reporter expression in SIX2^EGFP^ kidney organoids (Figure 3B and Figure 1C). As differentiation progressed, a steady increase in mCherry-expressing cells and a corresponding loss of EGFP-expressing cells was also observed (Figure 3B). Live mCherry^+^ cells could also be detected in kidney organoids by fluorescent microscopy, some of which appeared to be localized within EpCAM^+^ epithelial structures (Figure 3C). Whole-mount immunofluorescence of day 25 SIX2^Cre/Cre^:GAPDH^dual^ organoids was performed to determine the precise location of mCherry-expressing cells within specific cellular compartments, using markers specific to nephrons (WT1, NPHS1, LTL, ECAD, EpCAM), collecting duct (GATA3, ECAD), renal interstitium (MEIS1) and endothelium (CD31). *SIX2*-expressing cells contributed to nephron formation, as evidenced by the appearance of mCherry^+^ cells within LTL^+^ proximal tubules, ECAD^+^/LTL^-^ distal tubules and within NPHS1^+^ podocytes (Figure 3D). Consistent with our scRNAseq analysis, interstitial cells co-expressing MEIS1 and mCherry were also clearly evident, indicating that *SIX2*^+^ cells give rise to at least a subset of the renal stroma (Figure 3D). Conversely, CD31^+^/mCherry^+^ cells were not observed, indicating that endothelial cells within kidney organoids are not derived from *SIX2*^+^ cells (Figure 3D). Similarly, mCherry^+^ cells were excluded from the GATA3^+^/ECAD^+^ epithelium, suggesting that collecting duct cells within kidney organoids also arise from a population that is distinct from the *SIX2*^+^ population (Figure 3E). This was quantified and confirmed using image analysis software, where <1% of mCherry^+^ cells could be detected within GATA3^+^/ECAD^+^ epithelial structures (Figure 3F). Our findings are consistent with previous studies performed in mouse, which suggest the collecting duct network is instead derived from a more anterior ureteric progenitor population(Kobayashi et al., 2008; Taguchi et al., 2014) (Figure 3G).

**Figure 3.**
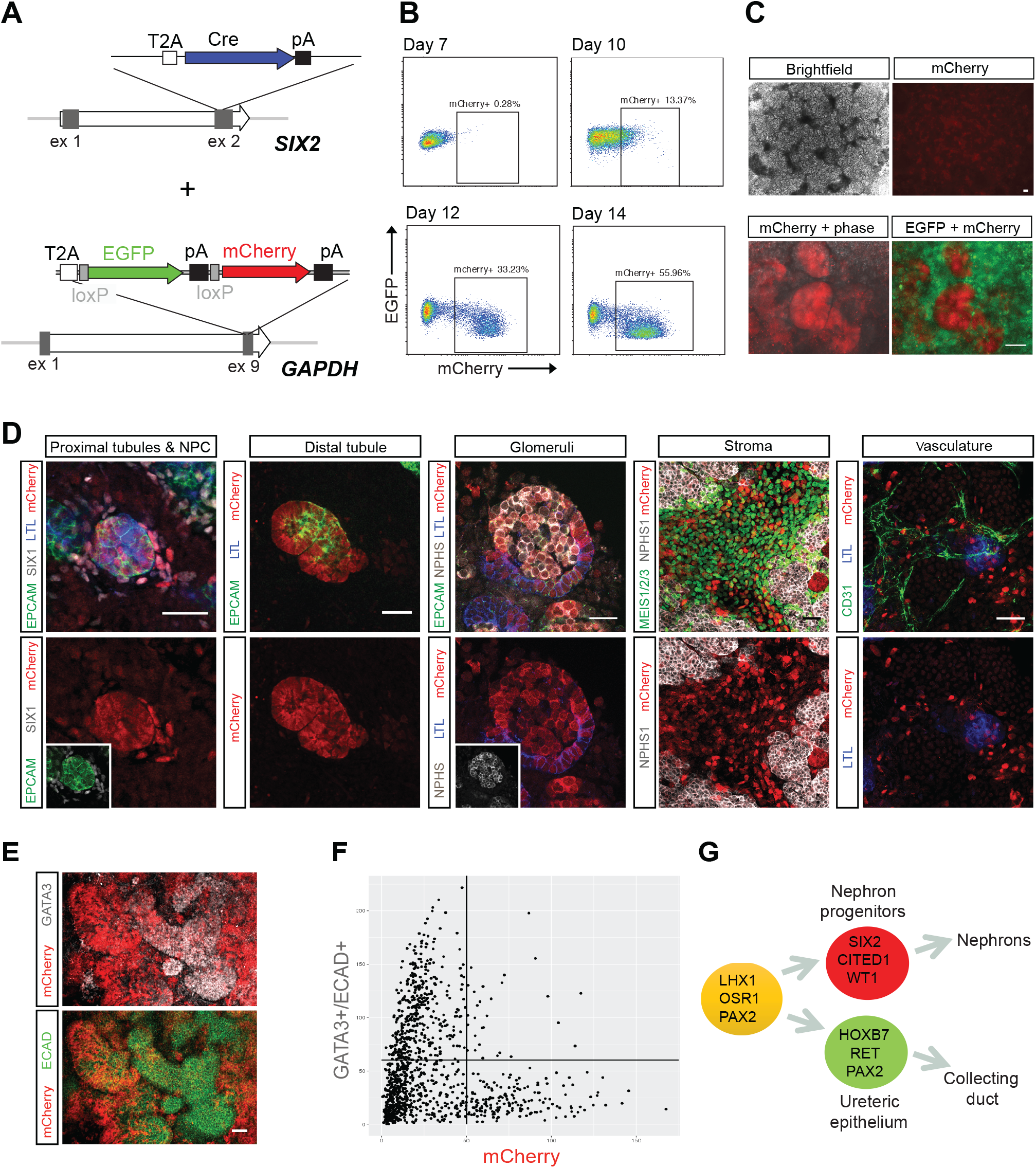
Fate mapping of the SIX2 population in hPSC-derived kidney organoids. (A) Schematic diagram of the targeting strategy used for generation of SIX2^Cre/Cre^:GAPDH^dual^ iPSCs. (B) Flow cytometry analysis of kidney organoids derived from SIX2^Cre/Cre^:GAPDH^dual^ iPSCs showing induction of mCherry and corresponding loss of EGFP expression. (C) Low (upper panel) and high magnification (lower panel) images showing mCherry^+^ cells detected by live fluorescent microscopy in SIX2^Cre/Cre^:GAPDH^dual^ kidney organoids. (D) Immunostaining of SIX2^Cre/Cre^:GAPDH^dual^ kidney organoids shows localization of mCherry cells within proximal (LTL^+^/EpCAM^+^), distal (LTL-/EpCAM^+^) and glomerular (NPHS1^+^) nephron segments and within renal stroma (MEIS1^+^) but not within the CD31^+^ vasculature. (E) mCherry^+^ cells were excluded from the presumptive GATA3^+^/ECAD^+^ collecting duct epithelium. (F) Plot showing exclusion of mCherry^+^ cells from GATA3^+^/ECAD^+^ epithelium as determined by image analysis software. (G) Model depicting the separation of nephron and collecting duct lineages during kidney morphogenesis. Scale bar = 50 μm.

### SIX2^+^ Cells Contribute to Nephron Formation in Early Stages of Organoid Differentiation

In the developing kidney *in vivo*, new nephrons arise throughout fetal development, arising from a self-renewing *Six2*^+^ nephron progenitor population present around the tips of the ureteric epithelium (Kobayashi et al., 2008). This process continues until approximately week 36 in human (Hinchliffe et al., 1991; Ryan et al., 2018) and the first few days after birth in mouse (Rumballe et al., 2011), at which point all NPC have committed to nephron formation. To determine the duration of nephrogenesis across the period of kidney organoid culture, we generated *SIX2* knock-in iPSCs using the tamoxifen-inducible Cre recombinase, CreERT2 (Indra et al., 1999). The dual fluorescence cassette was subsequently inserted into the endogenous *GAPDH* locus of a clonally derived iPSC line harboring a homozygous insertion of CreERT2 (Figure 4A). Day 12 kidney organoids derived from SIX2^CreERT2/CreERT2^:GAPDH^Dual^ iPSCs were cultured in the presence of 4-hydroxy-tamoxifen (4-OHT) for 1 hour to induce Cre recombination. Activation of the mCherry reporter could be detected by flow cytometry and fluorescent microscopy within 24 hours post-treatment (Figure 4B). A dose-dependent trend was also noted, with the number of mCherry^+^ cells positively correlating with 4-OHT concentration as anticipated (Figure 4C). Importantly, no mCherry^+^ cells were observed in the absence of 4-OHT treatment. To determine if *SIX2*^+^ cells could contribute to nephron formation throughout organoid development, we staggered the labeling of *SIX2*-expressing cells by initiating 4-OHT treatment (1 μM) at two day intervals between day 10–18 of differentiation (Figure 4D). Organoids were dissociated at day 25, stained with a directly-conjugated EpCAM antibody and analyzed by flow cytometry to determine the percentage of mCherry^+^ cells localized within epithelial structures. Using this assay, we observed significantly fewer EpCAM^+^ cells within the mCherry^+^ fraction of organoids induced at day 18 compared to those induced at day 10 (Figure 4E and Supplementary Figure 2A). While a negative correlation between the time of 4-OHT treatment and EpCAM^+^/mCherry^+^ cells was observed, there was no correlation between time of 4-OHT treatment and the total number of mCherry^+^ cells (Supplementary Figure 1B) nor the overall fraction of EpCAM^+^ cells (Supplementary Figure 1C). Whole-mount immunofluorescence of day 25 SIX2^CreERT2/CreERT2^:GAPDH^dual^ organoids was also performed, where we observed mCherry^+^ cells both within EpCAM^+^ structures and the interstitial compartment, but not within the GATA3^+^/EpCAM^+^ epithelium (Figure 4F). In organoids induced early, at day 10, mCherry^+^ cells were easily detected in all nephron epithelial segments and EpCAM^-^/NEPHRIN^+^ podocytes (Figure 4G, early induction). Conversely, in organoids induced at later time-points, mCherry^+^ cells were notably absent within nephron structures and predominantly restricted to the interstitium surrounding them (Figure 4G, late induction). Collectively, these findings indicate that the capacity for *SIX2*^+^ cells to contribute to nephron formation is largely restricted to the early stages of organoid differentiation and suggests that human kidney organoids lack a NPC niche that is capable of sustained nephrogenesis.

**Figure 4.**
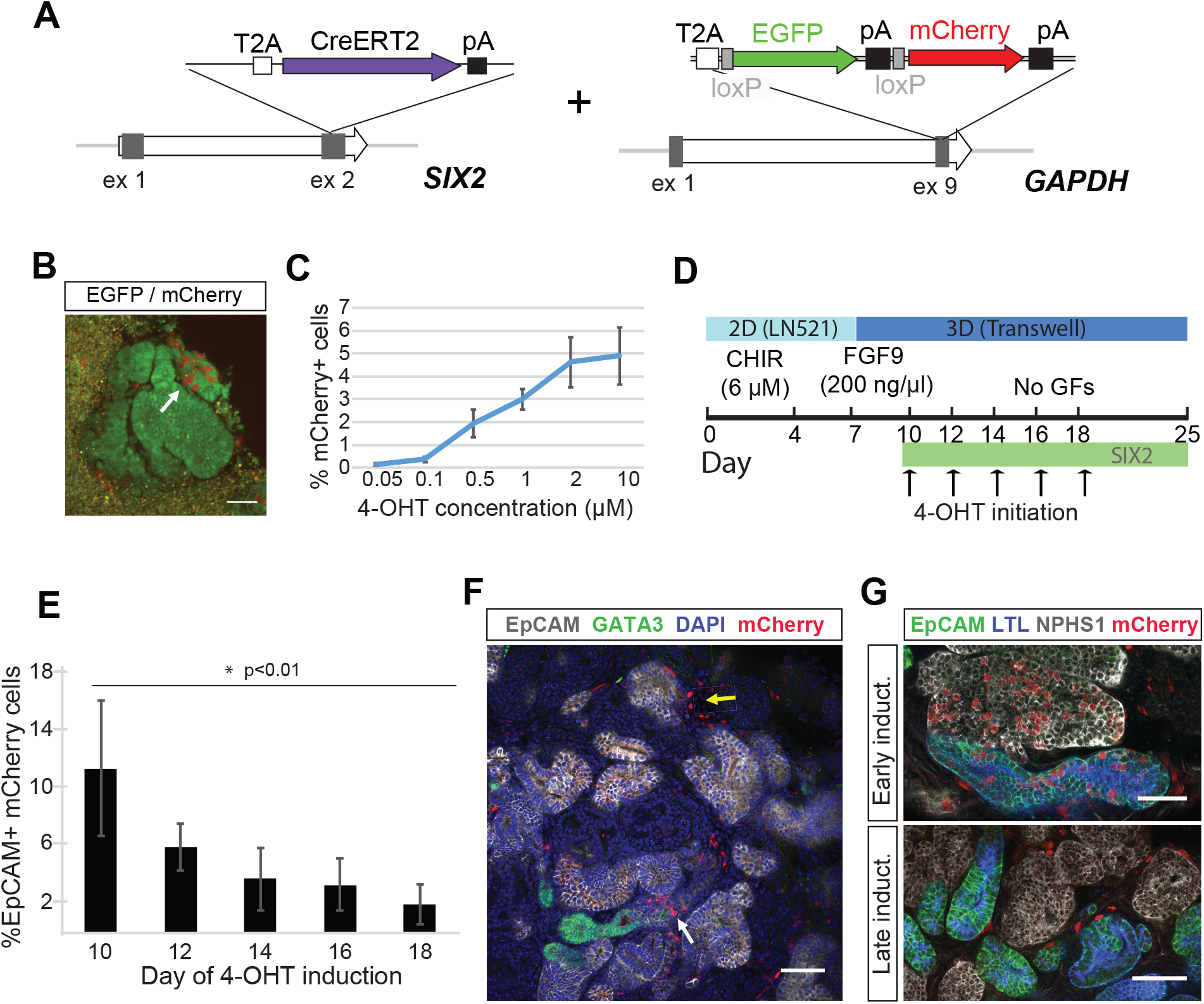
Temporal induction of SIX2-lineage shows declining nephrogenic potential of *SIX2*+ cells during kidney organoid differentiation. (A) Schematic diagram of the targeting strategy used for generation of SIX2^CreERT2/CreERT2^:GAPDH^dual^ iPSCs. (B) mCherry^+^ cells detected by live fluorescent microscopy in SIX2^CreERT2/CreERT2^:GAPDH^dual^ kidney organoid following 4-OHT induction. (C) The number of mCherry^+^ cells in kidney SIX2^CreERT2/CreERT2^:GAPDH^dual^ organoids positively correlates with 4-OHT concentration. Data represent mean ± SD, n=3. (D) Outline of the strategy used to interrogate the nephrogenic potential of *SIX2*^+^ cells throughout kidney organoid differentiation. (E) The percentage of mCherry^+^ cells localized within EpCAM^+^ epithelial structures negatively correlates with the time of 4-OHT initiation, as determined by flow cytometric analysis. Data represent mean ± SD, n≥3. Data from day 10 and 18 time-points was obtained from two independent experiments. See also Supplementary Figure 2. (F) Immunostaining of SIX2^CreERT2/CreERT2^:GAPDH^dual^ kidney organoids shows mCherry^+^ cells localized within EpCAM^+^ structures (white arrow) and the interstitial compartment (yellow arrow), but not within the GATA3^+^/EpCAM^+^ epithelium. (G) mCherry^+^ cells were detected in all nephron epithelial segments when 4-OHT-induction was initiated at day 10 (early induction) but were largely restricted to the interstitium when induced at later time-points (late induction). Scale bars = 50 μm.

## Discussion

Organoids derived from hPSCs offer enormous utility in personalized disease modeling and drug testing platforms, while also providing promise for the development of autologous cellular therapies to treat/correct many inherited and acquired diseases. Organoid-based cultures also represent a potential source of human tissue at developmental stages that are typically unavailable for research purposes. In combination with gene-editing technologies, this could facilitate the study of gene function and cellular process that govern human development *in vitro*. Genome engineering also facilitates studies aimed at characterizing the cell types produced in organoid-based cultures and how well these compare with primary developing tissue, which is imperative for understanding and addressing limitations associated with hPSC-derived tissue.

In this study, we use gene-edited iPSCs to interrogate the lineage relationships driving nephrogenesis within human kidney organoids to examine how this compares with our existing understanding of mammalian nephrogenesis *in vivo*, which is almost entirely based on studies performed in mouse. We used our previously described kidney organoid differentiation protocol and a newly derived *SIX2* iPSC reporter line to monitor *SIX2*^+^ NPCs in developing kidney organoids. *SIX2*^+^ cells emerged soon after organoid formation and, somewhat unexpectedly, persisted until the termination of differentiation at day 25. This is in direct contrast to other previously described differentiation protocols that report the rapid loss of *SIX2*^+^ NPCs soon after the formation of 3D kidney organoids (Morizane et al., 2015).

While single cell transcriptome profiling of organoids generated in this study revealed a distinct *SIX2*^+^ population that exhibited strong congruence with human fetal NPCs, *SIX2* expression was also detected in several additional cell populations. Interestingly, this included an “off-target” muscle-like population that was enriched in late but not early organoids. *SIX2* expression was also detected in several clusters identified as “renal stroma”. Consistent with this observation, fate-mapping experiments in human kidney organoids revealed an obvious *SIX2*^+^ lineage-derived MEIS1^+^ stromal population. This contrasts sharply with studies performed in mouse, where a strict lineage boundary between NPCs and the interstitial progenitor cells (IPCs) that give rise to the renal stroma has been shown to exist (Kobayashi et al., 2014; Levinson and Mendelsohn, 2003; Mugford et al., 2008). However, recent single cell RNAseq analysis of human fetal kidney has revealed substantial overlap between NPCs and IPCs, with co-expression of *SIX2, MEIS1* and *FOXD1* detected within human NPCs (Lindstrom et al., 2018a). Although difficult to determine definitively whether the co-expression of NPC and stromal markers is an artifact of our organoid culture system, the findings from this previous study suggest that this is indeed a true species difference. It is also possible that in kidney organoids the NPC/IPC lineage boundary is less well defined. A recent study in mouse suggests that Pax2 suppresses the transdifferentiation of NPCs into renal interstitial cell types (Naiman et al., 2017) and thus, perhaps even a slight downregulation of *PAX2* expression in organoid NPCs is sufficient to promote transition towards a more stromal cell fate.

Using a novel lineage tracing iPSC reporter line, we demonstrate that *SIX2*-expressing cells can indeed contribute to all segments of the developing nephron but are excluded from the distal ends of the GATA3^+^/ECAD^+^ epithelial structures, suggesting that the presumptive collecting duct network in kidney organoids arises from a population that is distinct from *SIX2*^+^ progenitors. Of note, not all cells within each nephron had undergone Cre-mediated color switching, with many nephrons comprised of both mCherry^+^ and EGFP^+^ cells. One explanation for this is that Cre-recombination may not have been complete. Alternatively, there may be a genuine contribution of both *SIX2* expressing and non-expressing cells within forming nephrons. More studies are required to investigate this further.

By using a tamoxifen-inducible variant of Cre recombinase to stagger the labelling of *SIX2*^+^ cells during organoid differentiation, we noted a significantly reduced capacity for these cells to contribute to nephron formation over time. This suggests human kidney organoids, in contrast to fetal kidney *in vivo*, lack a true nephrogenic zone capable of sustained nephrogenesis. In mouse, correct localization of NPCs and IPCs around the tips of the ureteric epithelium enables reciprocal inductive signals between these populations. This is required for continued branching of the ureteric epithelium and both NPC self-renewal and commitment, which drives organogenesis throughout fetal development(Combes et al., 2016; Little and McMahon, 2012). Although there is evidence of NPC commitment in our organoid cultures, there is no evidence of a branching ureteric epithelium nor a distinct domain of NPCs within a self-renewing niche, suggesting that appropriate spatial organization and/or reciprocal interactions between NPCs, ureteric epithelium, and possibly also the renal stroma, are deficient.

Perhaps a more logical strategy for generating kidney organoids with a sustainable nephrogenic niche would be to derive homogenous populations of NPCs, IPCs and ureteric epithelium in parallel which could be subsequently combined in 3D culture, in a spatial arrangement that more accurately depicts the developing organ *in vivo*. Indeed this strategy was partially recapitulated in a recent study which demonstrated a substantially improved higher-order structure in kidney organoids derived from mouse PSCs, and was highlighted by an impressive capacity for the ureteric epithelium to undergo several rounds of branching morphogenesis (Taguchi and Nishinakamura, 2017). However, while a human branching ureteric epithelium was generated, there was not a successful reciprocal interaction between this epithelium and surrounding presumptive human metanephric mesenchyme, highlighting our limited understanding of the exact conditions required to recapitulate a self-renewing NPC niche *in vitro*.

In conclusion, our findings provide proof-of-concept that gene-editing and organoid technologies can be combined to facilitate fate-mapping studies in differentiating hPSCs. This not only provides a unique opportunity to investigate lineage relationships in real time but also in a higher-throughput and more cost-effective manner compared with mammalian models. Additional fate-mapping analyses, such as those described here, may also provide deeper insight into human kidney development, further facilitating the development of more efficient and robust protocols to generate renal cell types for downstream applications. While applied to kidney in this instance, this approach can be used to gain a better understanding of the lineage relationships governing other developmental processes, particularly where multicellular and complex tissue-specific organoid differentiation protocols have been established.

## Materials & Methods

### Cell lines

Human foreskin fibroblasts (ATTC ID: CRL-2429) were cultured in DMEM (Thermo Fisher Scientific) supplemented with 15% fetal bovine serum (FBS) (Hyclone) and 1X MEM Non-Essential Amino Acids Solution (Thermo Fisher Scientific) at 37°C, 5% CO_2_ and 5% O_2_. All iPSC lines were maintained and expanded at 37°C, 5% CO_2_ and 5% O_2_ in Essential 8 medium (Thermo Fisher Scientific) on Matrigel-coated plates with daily media changes and passaged every 3–4 days with EDTA in 1X PBS as previously described (Chen et al., 2011). The genomic integrity of iPSCs were confirmed by molecular karyotyping using Infinium CoreExome-24 v1.1 SNP arrays (Illumina) and expression of common pluripotency markers (TRA-1–81, SSEA-4, CD9, OCT4) was confirmed by immunofluorescence and flow cytometry.

### Kidney organoid production

The day prior to differentiation, cells were dissociated with TrypLE (Thermo Fisher Scientific), counted using a hemocytometer and seeded onto Laminin521-coated 6-well plates at a density of 50 ×10^3^ cells per well in Essential 8 medium. Intermediate mesoderm induction was performed by culturing iPSCs in TeSR-E6 medium (Stem Cell Technologies) containing 4–8 μM CHIR99021 (R&D Systems) for four days. On day 4, cells were switched to TeSR-E6 medium supplemented with 200 ng/mL FGF9 (R&D Systems) and 1 μg/mL Heparin (Sigma Aldrich). On day 7, cells were dissociated with TrypLE, diluted 5-fold with TeSR-E6 medium, transferred to a 15 ml conical tube and centrifuged for 5 minutes at 300 × *g* to pellet cells. The supernatant was discarded, cells were resuspended in residual medium and transferred directly into a syringe for bioprinting. Syringes containing the cell paste were loaded onto a NovoGen MMX bioprinter, primed to ensure cell material was flowing, and user-defined aliquots (5,000–100,000 cells per organoid) were deposited on 0.4 μm Transwell polyester membranes in 6-well plates (Corning). Following bioprinting, organoids were cultured for 1 hour in the presence of 6 μM CHIR99021 in TeSR-E6 medium in the basolateral compartment and subsequently cultured until Day 12 in TeSR-E6 medium supplemented with 200 ng/mL FGF9 and 1 μg/mL Heparin. From Day 12 to Day 25, organoids were grown in TeSR-E6 media medium without supplementation. Unless otherwise stated, kidney organoids were cultured until harvest at Day 25. For induction of CreERT2 protein in kidney organoids derived from SIX2^CreERT2^ iPSCs, 4-hydroxytamoxifen (Sigma Aldrich) dissolved in ethanol at a concentration of 100 μM was diluted to working concentration in TeSR-E6. Induction media (1 ml) was pipetted under the transwell, and individual drops from a 20 μL pipette were carefully placed on top of the organoids on the filter to ensure complete coverage. Organoids were incubated at 37°C for 1 hour. Induction media was removed by washing 3 times with TeSR-E6 media every 10 minutes, and then returned to the media they were in prior to induction.

### Vector construction

The SIX2:EGFP vector (pDNR-SIX2:EGFP) carries a targeting cassette encoding the T2A peptide and EGFP gene flanked by ∼700 bp and ∼450 bp of sequence corresponding to sequence immediately upstream and downstream of the SIX2 stop codon respectively. Two gBlocks (Integrated DNA Technologies) encoding the targeting cassette were inserted into the *AatII* and *EcoRI* sites of the pDNR-Dual (Clontech) plasmid vector. The pDNR-SIX2:Cre and pDNR-SIX2:CreERT2 targeting vectors were generated as described above but substituting the Cre recombinase and CreERT2 recombinase genes for EGFP respectively. The GADPH targeting vector encoding the dual fluorescence cassette (pGAPTrap-loxEGFPloxCherry) was generated by inserting sequence encoding the T2A peptide, loxP-flanked EGFP gene with SV40 polyA signal and adjacent mCherry gene with SV40 polyA signal into the *SfiI* and *ClaI* sties of the pGAPTrap-mtagBFP2-IRESMuro plasmid vector (after removal of the mtagBFP2-IRESMuro cassette). A sgRNA plasmid specific to the 3’ end of the SIX2 coding region (pSMART-sgRNA-SIX2) was generated by annealing ODNs SIX2_sgRNA1a and SIX2_sgRNA1b followed by ligation into the *BbsI* sites of the pSMART-sgRNA vector (Howden et al., 2016). A sgRNA plasmid specific to the 3’ end of the GAPDH coding region (pSMART-sgRNA-GAPDH) was generated by annealing ODNs GAPDH_sgRNA1a and GAPDH_sgRNA1b followed by ligation into the *BbsI* sites of the pSMART-sgRNA vector. All plasmids were propagated in DH5-alpha E.Coli (BIOLINE) and prepared for transfection using a Plasmid Maxi kit (QIAGEN).

### Generation of knock-in iPSCs

All SIX2 knock-in iPSCs (EGFP, Cre, CreERT2) were derived from human foreskin fibroblasts (ATCC: CRL-2429) using a previously described protocol that combines reprogramming and gene-editing in one-step (Howden et al., 2018). Episomal reprogramming plasmids (pEP4E02SET2K, pEP4E02SEN2L, pEP4E02SEM2K and pSimple-miR302/367), in vitro transcribed mRNA encoding the SpCas9-Gem variant (Howden et al., 2016), the pSMART-sgRNA-SIX2 plasmid and either the pDNR-SIX2:EGFP, pDNR-SIX2:Cre or pDNR-SIX2:CreERT2 targeting vectors were introduced into fibroblast using the Neon transfection system as described below. In vitro transcribed mRNA encoding a truncated version of the EBNA1 protein was also included to enhance nuclear uptake of the reprogramming plasmids (Chen et al., 2011; Howden et al., 2006). Genomic DNA was isolated from resulting iPSCs using the DNeasy Blood & Tissue Kit (QIAGEN) in accordance with the manufacturer’s protocol and PCR analysis was performed using GoTaq Green Master Mix (Promega) to identify correctly targetd clones. ODNs SIX2F and EGFPR flank the 5’ recombination junction of SIX2:EGFP knock-in iPSCs, whereas SIX2F and CreR flank the 5’ recombination junction of SIX2:Cre and SIX2:CreERT2 knock-ins. ODNs SV40paF and SIX2R flank the 3’ recombination junction of SIX2:EGFP, SIX2:Cre and SIX2:CreERT2 knockin iPSCs. Heterozygous and homozygous clones were distinguished using ODNs SIX2F and SIX2R. For knock-in of the dual fluoresecnece cassette, the *GAPDH* targeting construct (pGAPTrap-loxEGFPloxCherry) was cotransfected with pSMART-sgRNA-GAPDH and mRNA encoding SpCas9-Gem into hPSCs using the Neon transfection system as described below. Correctly targeted (EGFP-expressing) clones were identified by fluorescent microscopy.

### In vitro transcription

Capped and polyadenylated in vitro transcribed mRNA encoding SpCas9-Gem protein (Howden et., 2016) was generated using the mMESSAGE mMACHINE T7 ULTRA transcription kit (Thermo Fisher Scientific) according to the manufacturer’s recommendations. Plasmid template was linearized with PmeI endonuclease prior to transcription. LiCl was used to precipitate mRNA before resuspension. A truncated version of the EBNA1 protein (Howden et al., 2006), used to facilitate uptake of reprogramming plasmids, was transcribed using the mMESSAGE mMACHINE SP6 transcription kit (Thermo Fisher Scientific) according to the manufacturer’s recommendations.

### Cell Transfection

Transfections were performed using the Neon Transfection System (Thermo Fisher Scientific). Human fibroblasts or iPSCs were harvested with TrypLE (Thermo Fisher) 2 days after passaging and resuspended in Buffer R at a final concentration of 1 × 10^7^ cells/ml. Electroporation was performed in a 100 μl tip using 1400 V, 20 ms, 2 pulses for human fibroblasts, or 1100 V, 30 ms, 1 pulse for human iPSCs. Following electroporation fibroblasts were transferred to 6-well Matrigel-coated plates containing DMEM+15% FBS and switched to reprogramming medium (TeSR-E7 + 100 μM sodium butyrate) after 3 days, with medium changes every other day. Electroporated human iPSCs were plated on 6-well Matrigel-coated plates containing Essential 8 medium with 5 μM Y-27632 (Tocris).

### Flow cytometry

Prior to analysis, single kidney organoids were dissociated with 0.2 ml of a 1:1 TrypLE/Accutase solution in 1.5 ml tubes at 37°C for 15–25 min, with occasional mixing (flicking) until large clumps were no longer clearly visible. 1 ml of HBBS supplemented with 2% FBS was added to the cells before passing through a 40 μM FACS tube cell strainer (Falcon). Flow cytometry was performed using a LSRFortessa Call Analyzer (BD Biosciences). Data acquisition and analysis was performed using FACsDiva (BD) and FlowLogic software (Inivai). Gating was performed on live cells based on forward and side scatter analysis.

### Whole Mount Immunofluorescence

Organoids were transferred to 48 well plates for fixation and immunofluorescence procedures. Fixation was performed using ice cold 2% paraformaldehyde (PFA; Sigma Aldrich) for 20 minutes followed by 15 minutes washing in three changes of phosphate-buffered saline (PBS). For immunofluorescence, blocking and antibody staining incubations were performed on a rocking platform for 3 hours at room temperature or overnight at 4°C, respectively. Blocking solution consisted of 10% donkey serum with 0.3% Triton-X-100 (TX-100; Sigma Aldrich) in PBS. Antibodies were diluted in 0.3% TX-100/PBS. Primary antibodies were detected with Alexa Fluor-conjugated fluorescent secondary antibodies (Invitrogen), diluted 1:500. Organoids were washed in at least 3 changes of PBS for a minimum of 1 hour following primary and secondary antibody incubations. Imaging was performed in glass-bottomed dishes (MatTek) with glycerol-submersion using either the Zeiss LSM 780 or Dragonfly Spinning Disk confocal microscopes.

### Single cell transcriptional profiling and data analysis

Organoids were dissociated as described above (for flow cytometry) and passed through a 40 μM FACS tube cell strainer. Following centrifugation at 300 *g* for 3 minutes, the supernatant was discarded and cells resuspended in 50 μl TeSR-E6 medium. Viability and cell number were assessed and samples were run across separate runs on a Chromium Chip Kit (10X Genomics). Libraries were prepared using Chromium Single Cell Library kit V2 (10X Genomics) and sequenced on an Illumina HiSeq with 100 bp paired-end reads. Cell Ranger (v1.3.1) was used to process and aggregate raw data from each of the samples returning a count matrix. Quality control and analysis was performed in R using the Seurat package (v2.3.1)(Butler et al., 2018). Cells with more than 125,000 UMIs, less than 500 genes expressed or more than 20% reads assigned to mitochondrial genes were filtered out. UMI counts, percentage of mitochondrial and ribosomal gene expression, and cell cycle phase identity were regressed out. Genes with less than two counts across the whole dataset were also filtered out. The final dataset had 5,365 cells and 22,105 identified genes. The two samples were merged using a canonical correlation analysis (CCA) using 1,429 genes with the highest dispersion present in both samples. The CCA subspaces were aligned and the first 25 principal components based on these genes were used to build a graph, which was clustered at a resolution of 1.6.

## Acknowledgements

The Murdoch Children’s Research Institute is supported by the Victorian Government’s Operational Infrastructure Support Program. The MCRI Gene editing facility is supported by the Stafford Fox Foundation. MHL is a Senior Principal Research Fellow of the National Health and Medical Research Council, Australia (GNT1136085). This work was supported by the National Institutes of Health, USA (DK107344–01) and the NHMRC (GNT1100970).

## Author Contributions

S.E.H. and M.H.L conceived the study and wrote the manuscript; S.E.H performed gene-editing experiments, S.E.H, J.M.V, S.B.W. and K.T performed kidney differentiation experiments, J.M.V, S.B.W. and K.T performed immunostaining, S.B.W. performed scRNAseq analysis. All authors assisted in manuscript preparation.

